# nihexporter: an R package for NIH funding data

**DOI:** 10.1101/033456

**Authors:** Jay Hesselberth, Erin Baschal, Nick Ellinwood, Ashley Pacheco, Sally Peach, Mary Sweet, Yuying Wang

## Abstract

1

**Motivation:** The National Institutes of Health (NIH) is the major source of federal funding for biomedical research in the United States. Analysis of past and current NIH funding can illustrate funding trends and identify productive research topics, but these analyses are conducted *ad hoc* by the institutes themselves and only provide a small glimpse of the available data. The NIH provides free access to funding data via NIH EXPORTER, but no tools have been developed to enable analysis of this data.

**Results:** We developed the `nihexporter R` package, which provides access to NIH EXPORTER data. We used the package to develop several analysis vignettes that show funding trends across NIH institutes over 15 years and highlight differences in how institutes change their funding profiles. Investigators and institutions can use the package to perform self-studies of their own NIH funding.

**Availability:** The `nihexporter R` package can be installed via github.

**Implementation:** The `nihexporter` package is implemented in the R Statistical Computing Environment.

**Contact:** Jay Hesselberth, University of Colorado School of Medicine

## 2 Introduction

The National Institutes of Health (NIH) is the major source federal funds for biomedical research in the United States. The NIH budget is approved by Congress each year, and in fiscal year 2014, the total NIH budget was 30.3 billion dollars (REF). The NIH is divided into 25 institutes, each with its own focus and mission. For example, National Cancer Institue (NCI) focuses on malignant diseases; the National Institute for Allergy and Immune Disease focuses on the immune system and transmissible diseasei; and the National Institute for General Medical Sciences focuses on basic research without a specific disease focus. Each institute negotiates with the NIH director (currently Francis Collins, MD, PhD) for its yearly budget, with budget institutes ranging from millions to several billion dollars.

The NIH provides funds through competetive grants written by internal and external investigators, and the funds associated with these grants can be divided into ‘direct’ and ‘indirect’ costs. **Direct costs** are funds that are given to an investigator (or group of investigators) to conduct their proposed research. These funds buy supplies for the experiments and pay the salaries of people to do the work.

By contrast, **indirect costs** are funds that are paid to institutions associated with investigators, and are used to “keep the lights on”: they pay for infrastructure costs. However, the “indirect cost recovery” (ICR) rate of each institution, the fraction of each award the institute receives, is congressionally mandated, and there is a wide range in ICR rates. Some of the highest ICR rates are close to 100%, meaning that for every dollar an investigator receives, the institutions receive an equal amount.

NIH funding is an investment strategy: the institutes invest money in specific research areas, hoping for future returns in the form of publications, patents and skilled trainees. As with any investment strategy, a periodic review can help rebalance the portfolio in order to maximize returns. Analysis of NIH funding data has been performed internally by the NIH, or by contracted third-parties. Several of these analyses have highlighted funding trends and suggested metrics to gauge the ‘return’ on the NIH ‘investment’. For example, “productivity” can be examined as a function of the number of publications produced by grants per dollar of “direct costs”.

## 3 Methods

We downloaded NIH funding data from the NIH EXPORTER website in comma-separated value (CSV) format and parsed these data into R data files that each contain specific information:

- `projects` has information about projects in each fiscal year, keyed by `project.num`
- `org_info` has information about project organizations
- `project_orgs` links applications to organizations
- `project_pis` links PI information to a project
- `publinks` table links PubMed IDs to `project.num`
- `publications` links project IDs to PubMed IDs
- `patents` links patent IDs to project IDs

See the documentation in the R package for more information about each table.

The package also has several precomputed variables and tables that enable quick and easy exploratory analysis:

- `nih.institutes`: Two-letter format for 27 NIH institutes
- `project_io`: This table contains pre-computed values for overall project cost `(project.cost)`, as well as the number of publications `(n.pubs)` and patents `(n.patents)` associated with each project.

NIH EXPORTER provides access to the total costs of each grant in each fiscal year, comprising both direct and indirect costs.

We developed several analyses using the `dplyr, ggplot2` and `knitr` packages in the RStudio environment.

## 4 Results and Discussion

### 4.1 Project costs

Let’s look at the costs of grants over time for a few institutes:

~~~
library(scales)
library(stringr)
library(dplyr)
library(ggplot2)
library(nihexporter)
~~~

~~~
select_inst <- c('GM', 'AI', 'CA')
cost.over.time <- projects %>%
   select(institute, fy.cost, fiscal.year) %>%
   filter(institute %in% select_inst) %>%
   group_by(fiscal.year, institute) %>%
   summarize(yearly.cost = sum(fy.cost, na.rm = TRUE))
cost.over.time %>%
   ggplot(aes(x = factor(fiscal.year),
                   y = yearly.cost / 1e9,
                   group = institute, color = institute)) + geom_line() +
  geom_point() +
  theme_bw() +
  theme(legend.position = 'bottom') +
  xlab('Fiscal year') +
  ylab('Project costs (dollars in billions)')
~~~

### 4.2 Funding distributions

#### 4.2.1 By Institution

Let’s look *where* the money is going. This example illustrates linking of the `project`, `org_info` and `project_orgs` tables.

~~~
institution_funding <- projects %>%
   filter(activity == 'R01') %>%
   left_join(project_orgs, by = 'application.id') %>%
   group_by(org.duns, fiscal.year) %>%
   summarise(total.dollars = sum(fy.cost, na.rm = TRUE)) %>%
   ungroup() %>%
   left_join(org_info, by = 'org.duns') %>%
   arrange(desc(total.dollars))
~~~

~~~
institution_funding %>%
   select(fiscal.year, org.name, org.duns, total.dollars) %>%
   mutate(total.award.millions = total.dollars / 1e6) %>%
   select(-total.dollars) %>%
   head(10) %>%
   knitr::kable(caption='R01 grant dollars awarded to specific institutions', col.names=c('Fiscal year', 'Org.', 'DUNS', 'Dollars (millions)'))
~~~

**Figure 1:**
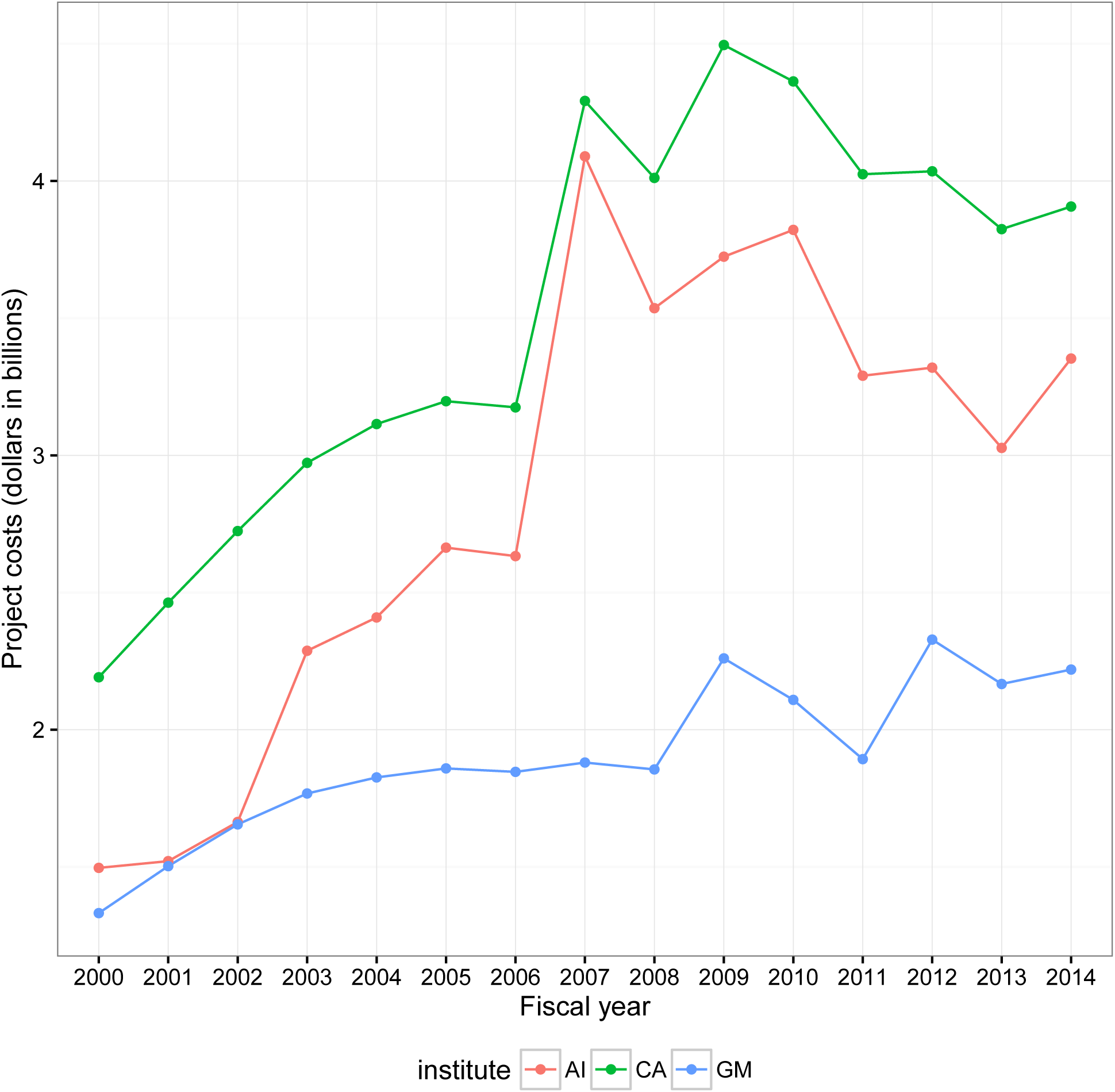
Project spending at NIGMS, NCI and NIAID

**Table 1:** R01 grant dollars awarded to specific institutions

| Fiscal year | Org. | DUNS | Dollars (millions) |
| --- | --- | --- | --- |
| 2009 | JOHNS HOPKINS UNIVERSITY | 1910777 | 297.2806 |
| 2010 | JOHNS HOPKINS UNIVERSITY | 1910777 | 292.9498 |
| 2004 | JOHNS HOPKINS UNIVERSITY | 1910777 | 289.2175 |
| 2011 | JOHNS HOPKINS UNIVERSITY | 1910777 | 286.3747 |
| 2005 | JOHNS HOPKINS UNIVERSITY | 1910777 | 278.1270 |
| 2012 | JOHNS HOPKINS UNIVERSITY | 1910777 | 273.9458 |
| 2006 | JOHNS HOPKINS UNIVERSITY | 1910777 | 270.8518 |
| 2007 | JOHNS HOPKINS UNIVERSITY | 1910777 | 268.4021 |
| 2003 | JOHNS HOPKINS UNIVERSITY | 1910777 | 266.1683 |
| 2010 | UNIVERSITY OF PENNSYLVANIA | 42250712 | 263.7944 |

We can also identify movers-and-shakers by institution.

#### 4.2.2 By PI

One can also examine how dollars are accrued by specific PIs. It is not possible to assign dollars directly to a PI, because some grants have multiple investigators. Rather, these are total costs that a given PI has been associated with over all grants in NIH EXPORTER. Here we identify PIs with the largest dollar amounts accrued for R01 grants.

~~~
dollars.per.pi <- projects %>%
   filter(activity == 'R01') %>%
   left_join(project_io, by = 'project.num') %>%
   left_join(project_pis, by = 'project.num') %>%
   filter(!is.na(pi.id)) %>%
   select(project.num, pi.id, total.cost) %>%
   unique() %>%
   group_by(pi.id) %>%
   summarise(pi.dollars = sum(total.cost) / 1e6) %>%
   arrange(desc(pi.dollars))
~~~

~~~
dollars.per.pi %>%
   head(10) %>%
   knitr::kable(digits = 3,
                     caption = 'R01 grant dollars associated with specific PIs',
                     col.names=c('PI ID', 'Dollars (millions)'))
~~~

**Table 2:**
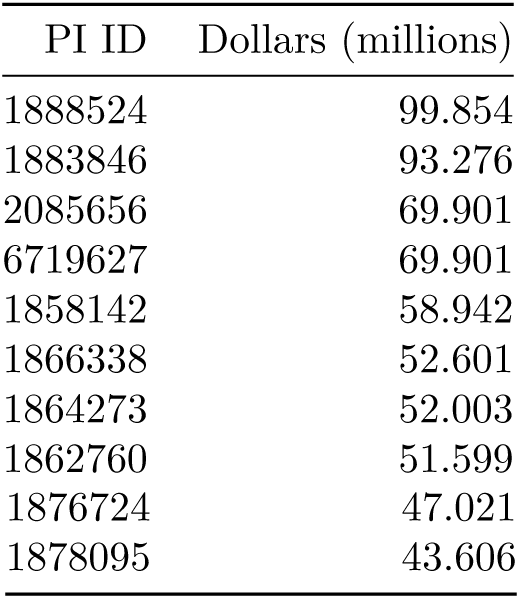
R01 grant dollars associated with specific PIs

### 4.3 Productivity

In order to measure the “return” on the money the NIH invests in the research enterprise, we can measure scholarly output (i.e., publications) per dollar invested.

Here we identify th highest performing grants outside of the R01 category. Much has been made of the wasteful spending outside of investigator-initiated research. Here we identify the cost of publications for grants other than R01’s that cost more than 1 million dollars.

~~~
high.perf.not.r01 <- projects %>%
   filter(activity != 'R01') %>%
   left_join(project_io, by = 'project.num') %>%
   select(project.num, total.cost, n.pubs) %>%
   filter(total.cost > 1e6 & n.pubs > 0) %>%
   mutate(cost.per.pub = total.cost / n.pubs) %>%
   arrange(cost.per.pub)
~~~

~~~
high.perf.not.r01 %>%
   head(10) %>%
   mutate(total.cost = round(total.cost / 1e6, 3),
            cost.per.pub = comma(round(cost.per.pub, 0))) %>%
   knitr::kable(caption='Productivity (publications / dollar) of non-R01 grants',
               col.names=c('Project ID', 'Project cost (dollars, millions)',
'Number of publications', 'Cost per publications (dollars)'))
~~~

**Table 3:** Productivity (publications / dollar) of non-R01 grants

| Project ID | Project cost (dollars, millions) | Number of publications | Cost per publications (dollars) |
| --- | --- | --- | --- |
| P30CA007175 | 2.449 | 721 | 3,397 |
| P30DK025295 | 2.565 | 691 | 3,712 |
| P30DK025295 | 2.565 | 691 | 3,712 |
| P30DK025295 | 2.565 | 691 | 3,712 |
| P30MH030929 | 1.710 | 444 | 3,851 |
| P30MH031302 | 1.046 | 266 | 3,931 |
| P01CA018221 | 2.727 | 692 | 3,941 |
| P01CA018221 | 2.727 | 692 | 3,941 |
| P41RR001008 | 1.506 | 366 | 4,115 |
| P41RR001008 | 1.506 | 366 | 4,115 |

We can also identify the specific publications associated with grants with the least expensive publications.

~~~
high.perf.not.r01 %>%
   head(1) %>%
   select(project.num) %>%
   left_join(publinks, by = 'project.num') %>%
   head(5) %>%
knitr::kable(caption='Publications from the most productive grants', col.names=c('Project ID', 'Pubmed ID'))
~~~

**Table 4:** Publications from the most productive grants

| Project ID | Pubmed ID |
| --- | --- |
| P30CA007175 | 3895298 |
| P30CA007175 | 4052630 |
| P30CA007175 | 3996611 |
| P30CA007175 | 2863007 |
| P30CA007175 | 4005847 |

We can also identify productive PIs with current R01s…

~~~
productive.pis <- projects %>%
   filter(activity == 'R01' & fiscal.year == 2014) %>%
   select(project.num) %>%
   left_join(project_io, by = 'project.num') %>%
   left_join(project_pis, by = 'project.num') %>%
   group_by(pi.id) %>%
   summarize(total.pi.dollars = sum(total.cost, na.rm = TRUE), total.pubs = sum(n.pubs)) %>%
   mutate(pub.cost = total.pi.dollars I total.pubs) %>%
   # *prevent PI Ids from being commafied* mutate(pi.id = as.character(pi.id)) %>%
   arrange(pub.cost)
~~~

~~~
productive.pis %>%
   head(10) %>%
   knitr::kable(digits = 0,
   format.args = list(big.mark = ','),
   col.names = c('PI ID',
                        'Cost per publication (dollars)',
                        'Total publications',
                        'Total project costs (dollars)'))
~~~

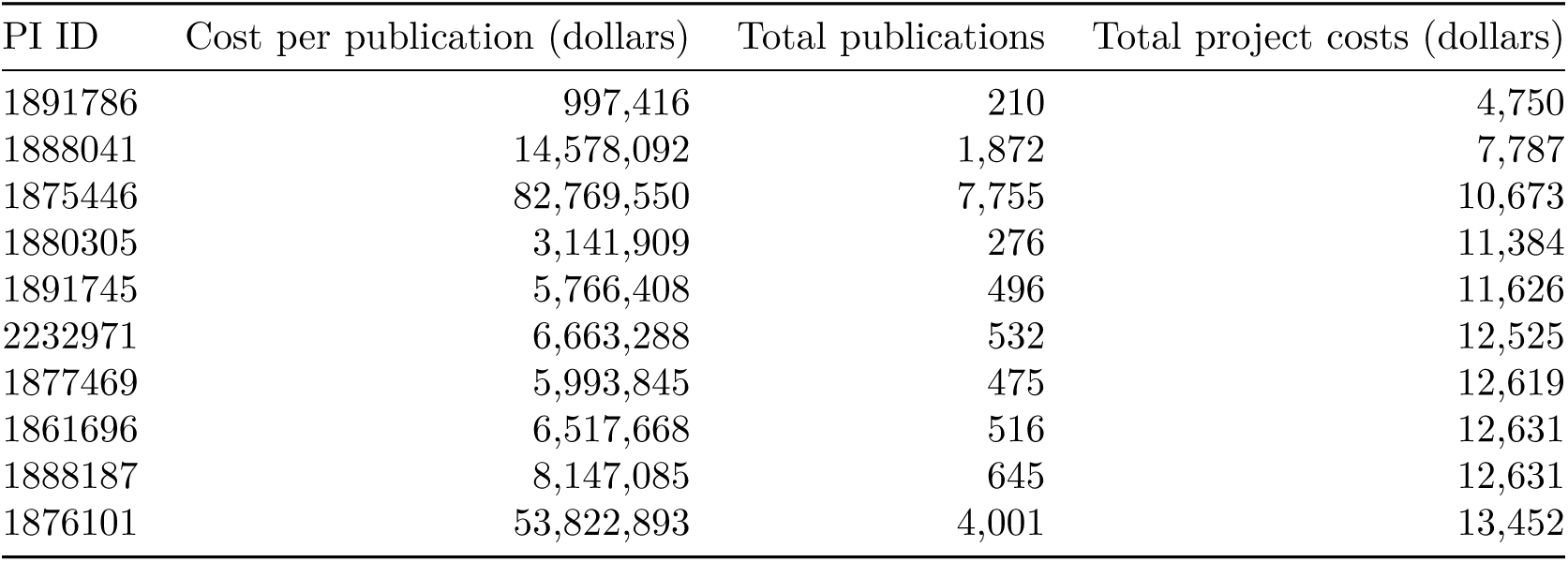

| PI ID | Cost per publication (dollars) | Total publications | Total project costs (dollars) |
| --- | --- | --- | --- |
| 1891786 | 997,416 | 210 | 4,750 |
| 1888041 | 14,578,092 | 1,872 | 7,787 |
| 1875446 | 82,769,550 | 7,755 | 10,673 |
| 1880305 | 3,141,909 | 276 | 11,384 |
| 1891745 | 5,766,408 | 496 | 11,626 |
| 2232971 | 6,663,288 | 532 | 12,525 |
| 1877469 | 5,993,845 | 475 | 12,619 |
| 1861696 | 6,517,668 | 516 | 12,631 |
| 1888187 | 8,147,085 | 645 | 12,631 |
| 1876101 | 53,822,893 | 4,001 | 13,452 |

Or we can identify the all-time most produtive PIs, independent of grant type or time frame…

~~~
productive.pis.all.time <- projects %>%
    select (project.num, activity) %>%
    left_join(project_io, by = 'project.num') %>%
    left_join(project_pis, by = 'project.num') %>%
    group_by(pi.id, activity) %>%
   summarize(total.pi.dollars = sum(total.cost, na.rm = TRUE),
                  total.pubs = sum(n.pubs),
                  total.patents = sum(n.patents)) %>%
    ungroup () %>%
   filter(total.pi.dollars >= 1e6) %>%
   mutate(pub.cost = total.pi.dollars / total.pubs,
             patent.cost = total.pi.dollars / total.patents) %>%
    select(pi.id, activity, pub.cost, patent.cost,
            total.pubs, total.patents, total.pi.dollars) %>%
   mutate(pi.id = as.character(pi.id))
~~~

~~~
productive.pis.all.time %>%
   select (-patent.cost, -total.patents) %>%
   arrange (pub.cost) %>%
   head(1O) 7.>7.
   knitr::kable(digits = 0,
                           format.args = list(big.mark = ','),
                           caption = 'Most productive PIs all time for publications',
                           col.names = c('PI ID', 'Grant type',
                                             'Cost per publication (dollars)',
                                             'Total publications',
                                             'Total dollars'))
~~~

**Table 6:** Most productive PIs all time for publications

| PI ID | Grant type | Cost per publication (dollars) | Total publications | Total dollars |
| --- | --- | --- | --- | --- |
| 1859859 | R03 | 716 | 3,184 | 2,278,288 |
| 1965193 | R01 | 1,065 | 3,024 | 3,221,984 |
| 1859703 | R01 | 1,143 | 5,257 | 6,009,650 |
| 7846601 | K01 | 1,191 | 10,400 | 12,387,225 |
| 2067600 | U10 | 1,452 | 1,000 | 1,451,976 |
| 7548099 | U10 | 1,452 | 1,000 | 1,451,976 |
| 1876003 | R37 | 1,659 | 738 | 1,224,549 |
| 2412990 | R13 | 1,996 | 9,200 | 18,366,900 |
| 6434316 | F32 | 2,154 | 558 | 1,202,148 |
| 1873876 | R37 | 2,339 | 657 | 1,536,750 |

~~~
productive.pis.all.time %>%
   select(-pub.cost, -total.pubs) %>%
   arrange(patent.cost) %>%
   head(10) %>%
   knitr::kable(digits = 0,
                    format.args = list(big.mark = ','),
                    caption = 'Most productive PIs all time for patents',
                    col.names = c('PI ID', 'Grant type',
                                       'Cost per patent (dollars)',
                                       'Total patents',
                                       'Total dollars'))
~~~

**Table 7:** Most productive PIs all time for patents

| PI ID | Grant type | Cost per patent (dollars) | Total patents | Total dollars |
| --- | --- | --- | --- | --- |
| 2081455 | R29 | 5,464 | 216 | 1,180,308 |
| 1900554 | R43 | 16,616 | 68 | 1,129,889 |
| 2104524 | R29 | 34,786 | 81 | 2,817,675 |
| 1883257 | R01 | 35,337 | 669 | 23,640,252 |
| 1863886 | R01 | 35,634 | 625 | 22,271,350 |
| 8913930 | R13 | 36,000 | 36 | 1,296,000 |
| 1866710 | R29 | 37,743 | 27 | 1,019,052 |
| 2415454 | T32 | 37,991 | 40 | 1,519,648 |
| 2185797 | R29 | 38,041 | 72 | 2,738,925 |
| 3129094 | R03 | 38,216 | 67 | 2,560,439 |

When Jeremy Berg was head of the Institute of General Medical Sciences (NIGMS) from 2003–2011, he routinely provided analysis of funding trends at NIGMS in his “Feedback Loop” blog. One of these measured the productivity per grant dollar by measuring its “scholarly output” *(i.e*., publications) as a function of direct costs. In this plot there is an increase in productivity per dollar, until an inflection point at $700K, after which the number of publications *drops*, suggesting a negative influence of grant money on scholarly output. This was interesting and covered here.

Here we flesh out this analysis and look at how all institutes perform by this measure (Berg, and now Lorsch, only analyzed GM). One caveat is that we only have access to total.cost in NIH EXPORTER, so the numbers include indirect costs. But, this is real cost to the tax-payer.

First, we need to calculate the lifetime costs of all R01 grants.

~~~
# *calculate costs of all grants, over the entire lifetime of the grant* grant_costs <- projects %>%
     filter(institute %in% nih.institutes & activity == 'R01') %>%
     left_join(project_io, by = 'project.num') %>%
     select(institute, project.num, total.cost, n.pubs) %>%
     unique()
~~~

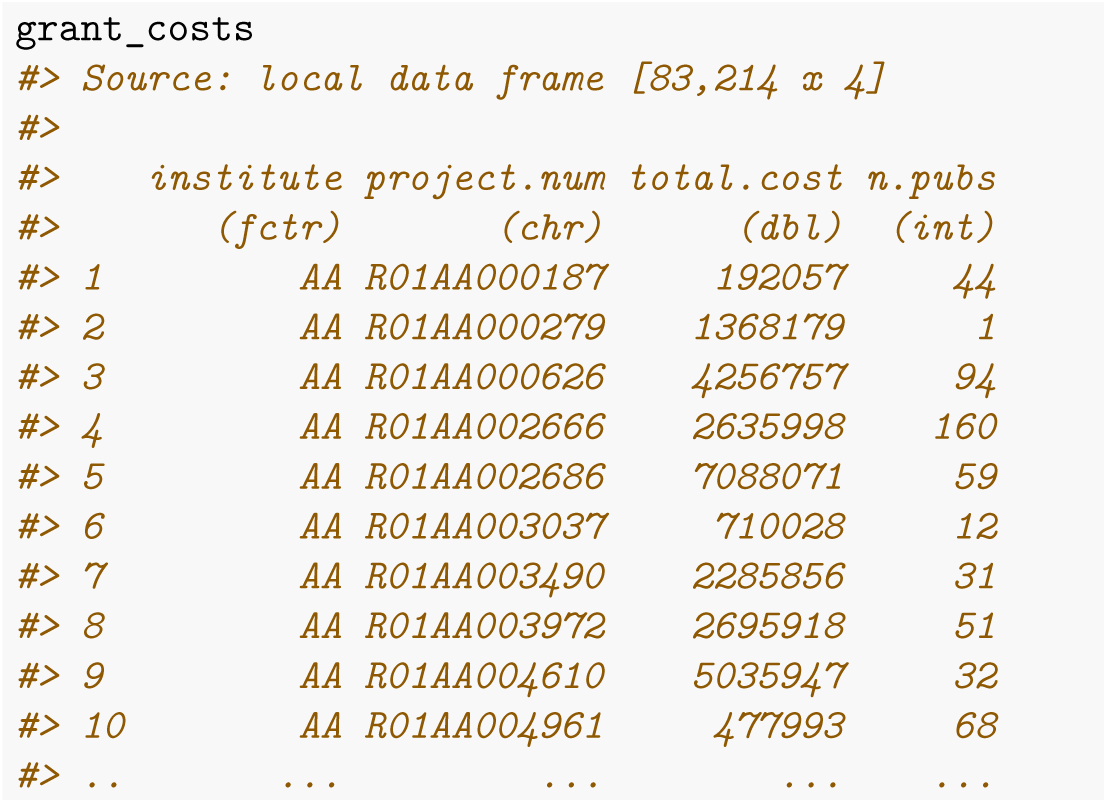

Next, we need to identify grants in each of the bins that Berg previously alluded to. `dplyr` makes this easy with the `ntile()` function. Berg previously divided grants into ~15 bins, we’ll bin into ~5%.

~~~
bin_grant_costs <- grant_costs %>%
   group_by(institute) %>%
   group_by(n.tile = ntile(total.cost, 20))
~~~

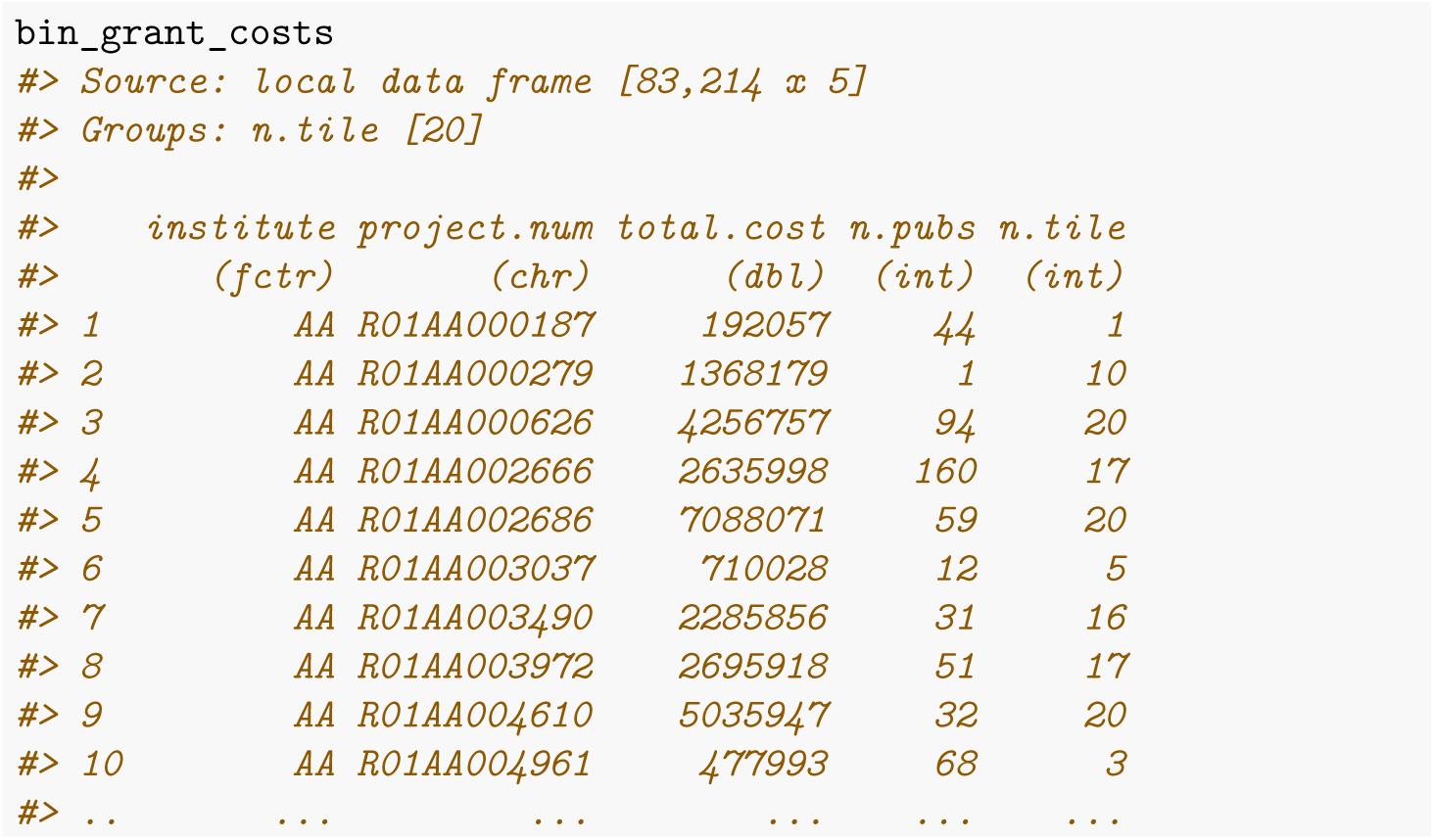

~~~
# *Berg's original values…*
# *breaks <- c(175000, 200000, 225000, 250000, 300000, 375000, 400000*,
#                  *450000, 500000, 600000, 700000, 800000, 900000, 1000000)*
min.lifetime.cost <- round(min(grant_costs$total.cost), -4) *# round to 10,000s* max.lifetime.cost <- round(max(grant_costs$total.cost), -5)
# *step is average size of an award* step <- 1e6
~~~

~~~
breaks <- seq(min.lifetime.cost, max.lifetime.cost, step) breaks
#> [1] 0.0e+00 1.0e+06 2.0e+06 3.0e+06 4.0e+06 5.0e+06 6.0e+06 7.0e+06
#> [9] 8.0e+06 9.0e+06 1.0e+07 1.1e+07 1.2e+07 1.3e+07 1.4e+07 1.5e+07
#> [17] 1.6e+07 1.7e+07 1.8e+07 1.9e+07 2.0e+07 2.1e+07 2.2e+07 2.3e+07
#> [25] 2.4e+07 2.5e+07 2.6e+07 2.7e+07 2.8e+07 2.9e+07 3.0e+07 3.1e+07
#> [33] 3.2e+07 3.3e+07 3.4e+07 3.5e+07 3.6e+07 3.7e+07 3.8e+07 3.9e+07
#> [41] 4.0e+07 4.1e+07 4.2e+07 4.3e+07 4.4e+07 4.5e+07 4.6e+07 4.7e+07
#> [49] 4.8e+07 4.9e+07 5.0e+07 5.1e+07 5.2e+07 5.3e+07 5.4e+07 5.5e+07
#> [57] 5.6e+07 5.7e+07 5.8e+07 5.9e+07 6.0e+07 6.1e+07 6.2e+07 6.3e+07
#> [65] 6.4e+07 6.5e+07 6.6e+07 6.7e+07 6.8e+07 6.9e+07 7.0e+07 7.1e+07
#> [73] 7.2e+07 7.3e+07 7.4e+07 7.5e+07 7.6e+07 7.7e+07
~~~

~~~
dollar_bin_grant_costs <- grant_costs %>%
   group_by(institute) %>%
   mutate(dollar.tile = findInterval(total.cost, vec = breaks, all.inside = TRUE, rightmost.closed = TRUE))
~~~

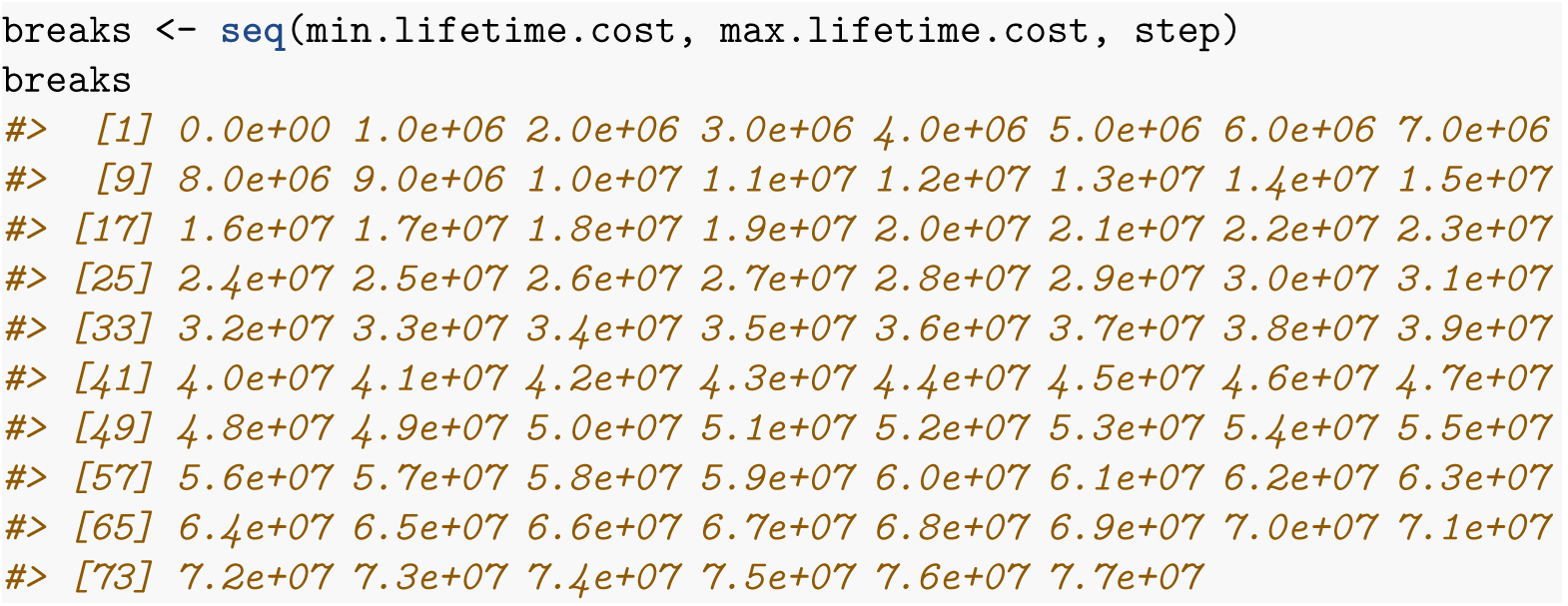

dollar_bin_grant_costs %>% group_by(dollar.tile) %>% summarize(count = n())

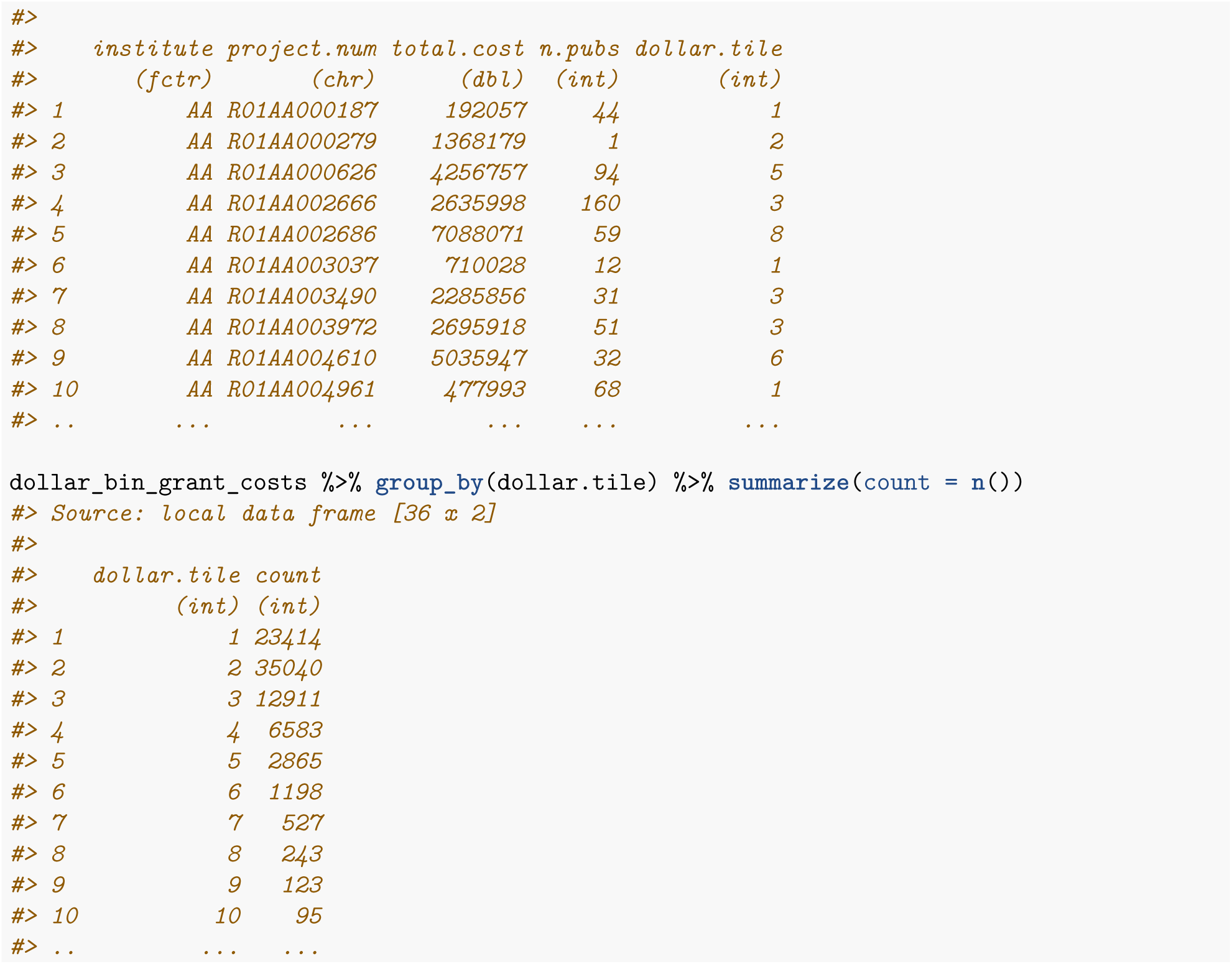

That looks better. Now we can make the summary plots…

~~~
dollar_bin_grant_costs %>%
   # *need to remove higher tiles because there are too few grants*
   filter(dollar.tile <= 13) %>%
   ggplot(aes(x = factor(dollar.tile), y = n.pubs)) +
   geom_boxplot(outlier.shape = NA) +
   scale_x_discrete(labels = breaks / 1e6) +
   theme(axis.text.x = element_text(angle=45, vjust=0.8)) +
   scale_y_log10() +
   theme_bw() +
   facet_wrap(~ institute, scales = "free_x") +
   ylab('Number of publications') +
   xlab('Total costs (minimum, in millions)')
~~~

### 4.4 Comparison of grant programs

The NIH provides funds through differet grant programs:

- `research`: investigator-intitiated, actvities begin with R (e.g., R01)
- `program`: activities begin with P (e.g. P01)
- `contract`: actvities begin with U (e.g. U54)

**Figure 2:**
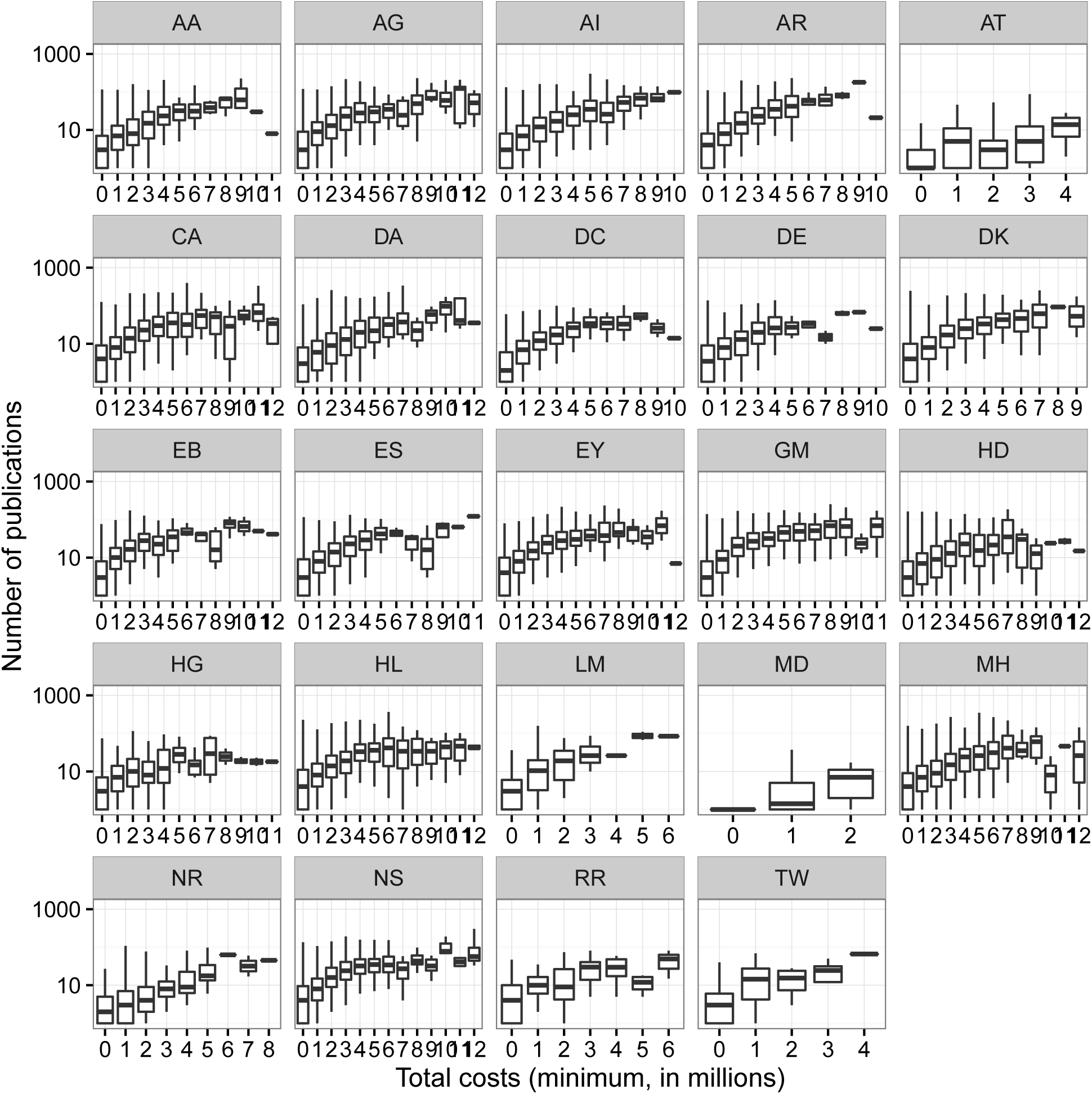
Productivity versus grant costs for each institute.

We can examine the total costs spent on specific grants and specific institutes over time.

**Figure 3:**
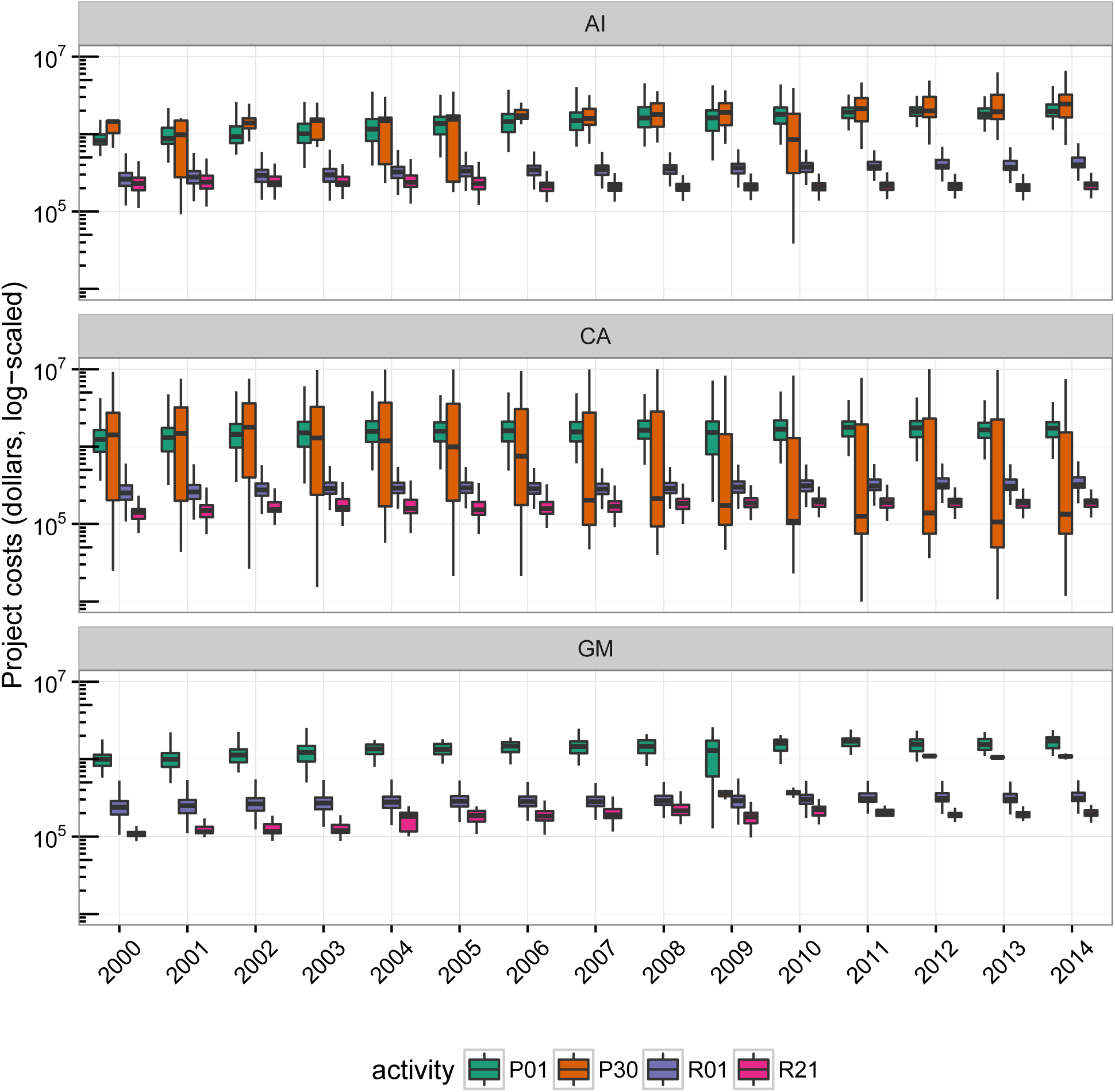
Funds for specific types of grants at the GM, CA and AI institutes

Or we can see how institutes allocate money generally to different types of grants.

~~~
# *from http://grants.nih.gov/grants/funding/funding_program.htm*
research_projects <- projects %>% filter (grepl(' ̑R', activity)) %>% select (project.num)
program_projects <- projects %>% filter (grepl ('̑P ', activity)) %>% select (project.num)
contract_projects <- projects %>% filter (grepl ('̑U', activity)) %>% select (project.num)
select_inst <- c('AI','CA','GM','HG','AA','MH')
~~~

~~~
grant_costs <- projects %>%
   filter (institute %>%    nih.institutes) %>%
    select(project.num, institute, fiscal.year, fy.cost)
~~~

~~~
research_costs <- grant_costs %>%
   semi_join(research_projects, by = 'project.num') %>%
    group_by(project.num, institute, fiscal.year) %>%
    summarize(project.cost = sum(fy.cost, na.rm = TRUE)) %>%
    mutate(type = 'research')
~~~

~~~
program_costs <- grant_costs %>%
   semi_join(program_projects, by = 'project.num') %>%
    group_by(project.num, institute, fiscal.year) %>%
    summarize (project.cost = sum(fy.cost, na.rm = TRUE)) %>%
    mutate(type = 'program')
~~~

~~~
contract_costs <- grant_costs %>%
   semi_join(contract_projects, by = 'project.num') %>%
    group_by(project.num, institute, fiscal.year) %>%
    summarize (project.cost = sum(fy.cost, na.rm = TRUE)) %>%
    mutate(type = 'contract')
~~~

~~~
combined_tbl <- bind_rows(research_costs, program_costs, contract_costs)
ggplot(combined_tbl, aes(x = factor(fiscal.year),
                                      y = project.cost,
                                         fill = type)) +
   geom_boxplot(outlier.shape = NA) +
   scale_y_log10(limits = c(1e4,1e7),
                         labels = trans_format("log10", math_format(10̑.x))) +
   facet_wrap(-institute) +
   theme_bw() +
   scale_fill_brewer(palette = "Dark2") +
   theme(legend.position = 'bottom') +
   xlab('') +
   ylab('Total grant costs, log-scaled') +
   theme(axis.text.x = element_text(angle = 90))
~~~

### 4.5 Duration

The `nihexporter` exposes `project.start` and `project.end`, which we can use to examine the duration of projects. For example, we can identify the longest running R01 grants.

~~~
long.grants <- projects %>%
   filter (activity == 'R01') %>%
   select (project.num, project.start, project.end) %>%
    group_by(project.num) %>%
   summarize (longest.run = max (project.end) - min(project.start)) %>%
    arrange(desc(longest.run)) %>%
   mutate(in.years = as.numeric(longest.run) I 365) %>%
   select(project.num, in.years)
long.grants %>%
   head(10) %>%
   knitr::kable(digits = 2,
                    caption = 'Longest running R01 grants (all-time)',
                    col.names=c('Project ID', 'Duration (years)'))
~~~

**Figure 4:**
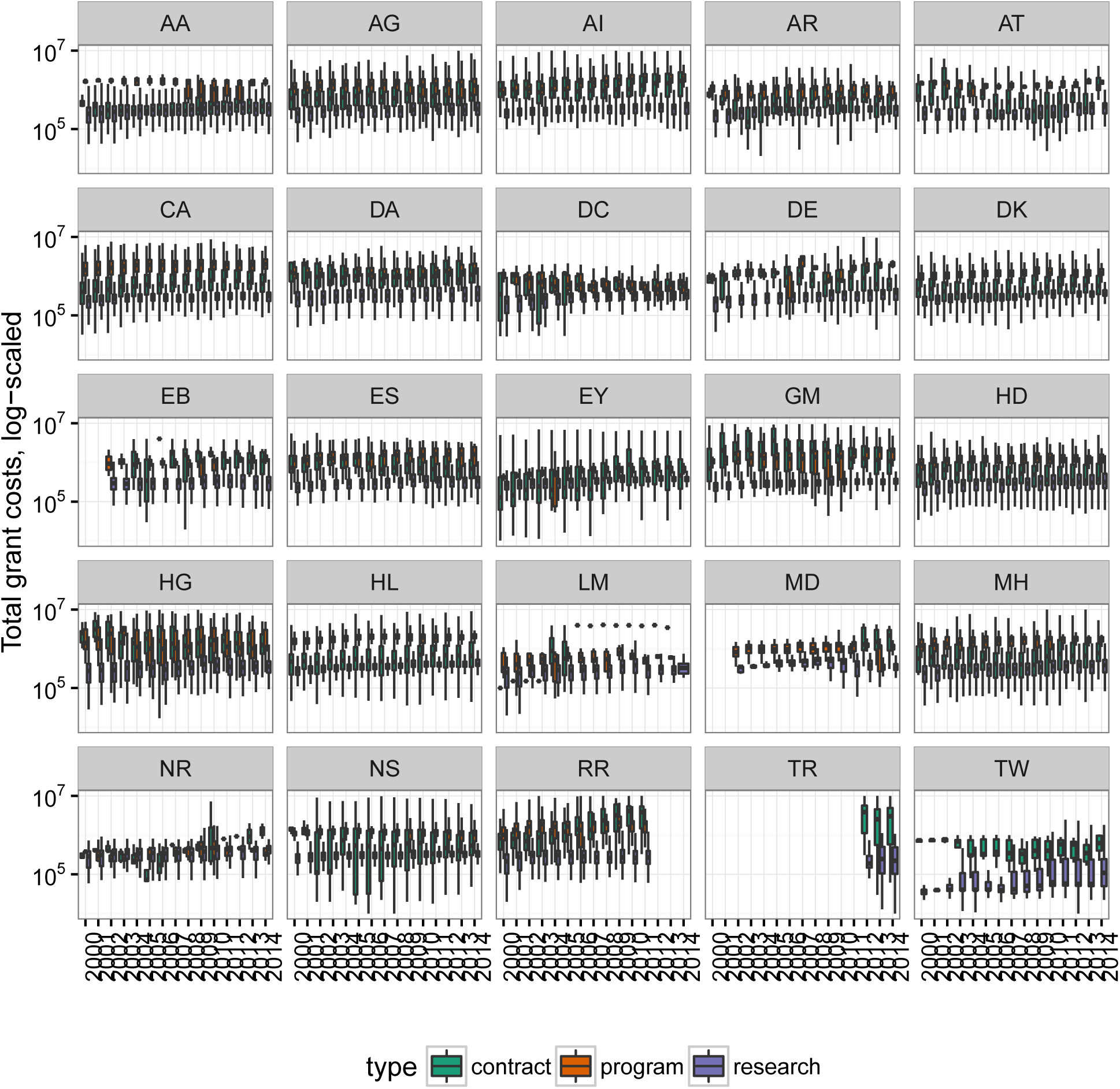
Comparison of research, program and contract costs over time

**Table 8:** Longest running R01 grants (all-time)

| Project ID | Duration (years) |
| --- | --- |
| R01HL009610 | 50.03 |
| R01HL013629 | 47.03 |
| R01HD008188 | 44.61 |
| R01CA013202 | 42.86 |
| R01EB001960 | 42.19 |
| R01DA001411 | 41.87 |
| R01GM023303 | 41.70 |
| R01HL017964 | 41.52 |
| R01DC000117 | 41.44 |
| R01EY002686 | 41.28 |

#### 4.5.1 Geographical distribution

Geographical distribution of grant dollar is easily visualized using the package.

~~~
state.data <- data.frame(org.state = state.abb,
                                   state.name = tolower(state.name))
~~~

~~~
state_funding <- projects %>%
   select(application.id) %>%
   left_join(project_orgs) %>%
   left_join(org_info) %>%
   left_join(projects, by = 'application.id') %>%
   select(application.id, org.state, fy.cost) %>%
   group_by(org.state) %>%
   summarize(total.fy.cost = sum(fy.cost) / 1e9)
~~~

~~~
cost.by.state <- state_funding %>%
   left_join(state.data) %>%
   select(state.name, total.fy.cost) %>%
   filter(state.name != "NA") %>%
   mutate(region=state.name, cost = total.fy.cost) %>%
   select(region, cost) %>%
   data.frame()
state.map.data <- map_data("state")
plot_data <- left_join(state.map.data, cost.by.state)
ggplot() +
geom_polygon(data=plot_data, aes(x=long, y=lat, group = group, fill=plot_data$cost), colour="black") +
scale_fill_continuous(low = "lightgrey", high = "red", guide="colorbar") +
theme_bw() +
labs(fill = "Total Cost per year \n (Billions of dollars)",x="", y="") + *#title = "Total cost per fiscal year by state"*
scale_y_continuous(breaks=c()) +
scale_x_continuous(breaks=c()) +
theme(panel.border = element_blank()) +
coord_fixed(ratio = 1.4)
~~~

**Figure 5:**
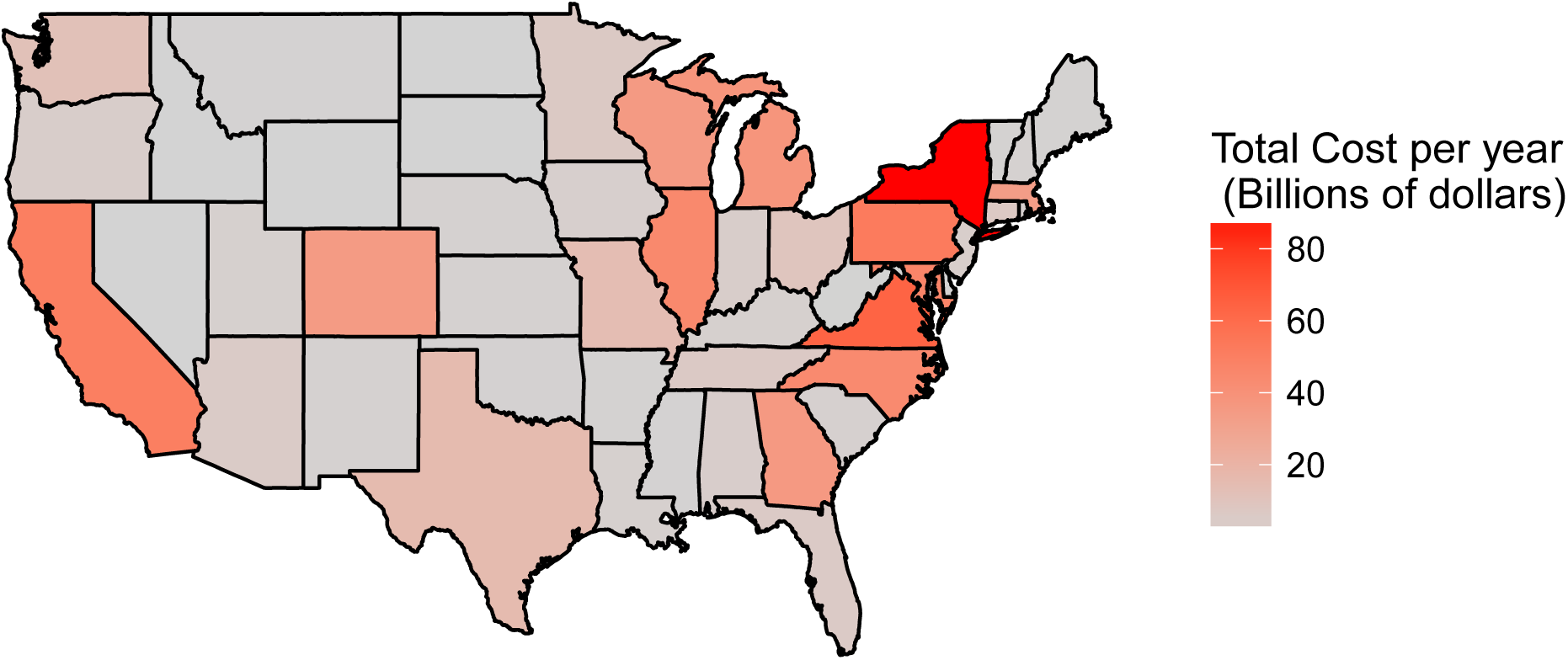
Geographic distribution of NIH dollars

## 5 Acknowledgments

The `nihexporter` package was developed over several weeks during a course on data analysis (http:// molb7621.github.io). Thanks to the NIH EXPORTER team for fixing issues with the EXPORTER tables.

